# MHC-Fine: Fine-tuned AlphaFold for Precise MHC-Peptide Complex Prediction

**DOI:** 10.1101/2023.11.29.569310

**Authors:** Ernest Glukhov, Dmytro Kalitin, Darya Stepanenko, Yimin Zhu, Thu Nguyen, George Jones, Carlos Simmerling, Julie C. Mitchell, Sandor Vajda, Ken A. Dill, Dzmitry Padhorny, Dima Kozakov

## Abstract

The precise prediction of Major Histocompatibility Complex (MHC)-peptide complex structures is pivotal for understanding cellular immune responses and advancing vaccine design. In this study, we enhanced AlphaFold’s capabilities by fine-tuning it with a specialized dataset comprised by exclusively high-resolution MHC-peptide crystal structures. This tailored approach aimed to address the generalist nature of AlphaFold’s original training, which, while broad-ranging, lacked the granularity necessary for the high-precision demands of MHC-peptide interaction prediction. A comparative analysis was conducted against the homology-modeling-based method Pandora [13], as well as the AlphaFold multimer model [8]. Our results demonstrate that our fine-tuned model outperforms both in terms of RMSD (median value is 0.65 Å) but also provides enhanced predicted lDDT scores, offering a more reliable assessment of the predicted structures. These advances have substantial implications for computational immunology, potentially accelerating the development of novel therapeutics and vaccines by providing a more precise computational lens through which to view MHC-peptide interactions.

## Introduction

MHC Class I (MHC I) molecules play a crucial role in the immune system and are found on the surface of most cells in the body. They present intracellular specific antigens, such as viral, bacterial, or cancerous peptides, to cytotoxic T cells, enabling T cells to recognize and respond to these threats [19].

MHC I molecules are important to the functioning of the immune system. By understanding how they bind and present peptides, we can gain insights into disease mechanisms such as autoimmunity [5]. This knowledge would also empower us to prevent certain infectious diseases through MHC-based vaccination [24]. In the context of cancer immunotherapy, this understanding would allow the design of neoantigen vaccines that enhance the immune system’s ability to selectively target cancer cells [20].

To ensure that the immune system effectively detects and responds to a wide range of infections, each MHC I molecule presents a variety of peptides to T cells. To achieve this, each MHC I has the capability to bind a broad class of different peptide sequences. Although each person presents only a small number of different MHC I molecules (two alleles for each of the three MHC I genes), many different MHC I alleles are present in the population [23], leading to individual differences in MHC I specificity. The diversity of MHC I molecules and peptides allows the immune system to respond effectively to different threats and adapt to new challenges. However, this diversity also poses a significant challenge when it comes to predicting MHC-peptide complexes (pMHC I).

There are different approaches for predicting pMHC I complex structures, including molecular docking [14], molecular dynamics simulations [18, 9], homology modelling [1] and machine learning methods [6, 15, 4]. Their accuracy of predictions can vary.

One of the advanced tools for predicting pMHC I complex structures is Peptide ANchoreD mOdelling fRAmework (PANDORA) [13]. Pandora uses a database of known MHC structures as templates with anchors-restrained loop modeling for peptide conformation. However, Pandora has some limitations: rare alleles may lack suitable MHC templates, low sequence similarity in the peptide-binding groove can cause alignment issues, accurate anchor residues are needed, and ranking output models can be challenging due to different scoring functions.

Transitioning from traditional homology-modeling techniques, AlphaFold introduces a revolutionary approach to protein structure prediction, utilizing deep learning to predict protein structures with remarkable accuracy. Its approach leverages a wealth of structure data to train its algorithm, enabling it to predict protein structures even in the absence of closely related known structures. However, despite AlphaFold’s successes, its utility for MHC-peptide predictions has been somewhat limited by its generalized training across diverse protein types. This broad approach, while comprehensive, may not always capture the intricate nuances necessary for high-fidelity predictions within specific domains such as MHC-peptide interactions. Therefore, by fine-tuning AlphaFold with a dataset curated explicitly from high-resolution MHC-peptide crystal structures, we aim to enhance the model’s specificity and accuracy in this critical area, thereby overcoming one of the main limitations faced by practitioners utilizing this tool for specialized applications in immunological research.

We present an approach that leverages the robust capabilities of AlphaFold, fine-tuning it to significantly improve the prediction accuracy for MHC-peptide complex structures. Our model outperforms Pandora in terms of CA RMSD (median value is 0.65 Å) and also provides enhanced predicted lDDT scores, offering a more reliable assessment of the predicted structures. Our model also does not require any input information about the anchor residues.

## Related Work

### pMHC I Structural overview

MHC I is a transmembrane protein composed of two non-covalently connected chains, *α* and *β*_2_ microglobulin [11]. The *α* chain consists of three domains *α*1, *α*2, *α*3 followed by transmembrane part and a cytoplasmic tail. Each *α* domain is approximately 90 residues long.

*α*1 and *α*2 domains form a symmetric structure composed of curved *α*-helices. Together they form a peptide-binding groove between them. The most variability in MHC I sequence is found in this groove region to create a variety for peptide specificity. *α*3 and *β*_2_ microglobulin domains do not interact with the peptide.

The number of different peptides that a particular MHC I molecule can bind varies depending on the specific MHC molecule. But in general peptide-binding groove geometry can accommodate short peptides of length 8–11 residues [16]. However, slightly longer peptides were also observed [22, 25, 10]. Several peptide positions, typically at the N and C termini of the peptide, contribute significantly to the binding. These specific residues are known as anchors.

### Homology modeling (Pandora)

One of the state-of-the-art tools for pMHC I complex structure prediction is the homology-based Peptide ANchoreD mOdelling fRAmework (PANDORA) [13]. Pandora uses homology modeling for the MHC protein and performs anchor-restrained peptide modelling using MODELLER.

For the homology modelling of MHC I, Pandora selects a single template from the custom-made database of known MHC structures. This approach requires proper template selection, which depends on the availability of the same allele type, group or gene within the database.

After selecting the template, the alignment between the target MHC and the template is carried out. The sequence similarity between the target and the template may be low in the peptide-binding groove due to MHC sequence variability, which can increase the likelihood of alignment issues. However, this groove region is of utmost importance for modeling since it is the area where peptide binds to MHC I.

One of the crucial requirements for Pandora is the inclusion of peptide anchors. Pandora utilizes this information for anchors-restrained loop modeling of the peptide. This information can be provided by the user or predicted by netMHCpan 4.1, although challenges may arise for non-canonical anchors.

Pandora provides 20 models for a single peptide-MHC pair and evaluates them using MODELLER’s internal scoring functions: molpdf and DOPE. In some cases, the best-scored DOPE and molpdf structures are different, which can present a challenging decision for the user in selecting the best model. The authors suggest using molpdf scoring ranking.

### AlphaFold

AlphaFold has been a transformative force in computational biology, redefining the landscape of protein structure prediction with its transformer-based architecture. This architecture is particularly well-suited to processing sequential biological data, such as amino acid chains, due to its ability to capture long-distance interactions between amino acids—a fundamental factor in predicting how proteins fold.

The model’s capabilities were brought to the forefront during the Critical Assessment of Structure Prediction (CASP) competitions, with AlphaFold setting new benchmarks in CASP13 and CASP14. Its exemplary performance highlighted the model’s broad potential, which extends beyond structure prediction to facilitating our understanding of disease pathology, advancing drug discovery, and innovating enzyme engineering.

However, the generalist nature of AlphaFold, while powerful, reveals limitations when tasked with highly specialized predictions, such as modeling the MHC-peptide complexes. The complexity inherent to these biological structures, coupled with their variability, calls for a tailored version of the model.

In the pursuit of enhancing AlphaFold’s predictions for MHC-peptide complexes, researchers have embarked on a variety of methodologies. One prominent approach involves the development of an AlphaFold-based pipeline which involves additional steps for MSA or template selection [15]. In addition, efforts have been made to fine-tune AlphaFold’s parameters on peptide-MHC Class I and II structural and binding data, with the fine-tuned model achieving state-of-the-art classification accuracy [17]. These advancements, while in some cases requiring more complex pipelines, reflect the nuanced balance between achieving broad predictive capabilities and the pursuit of granular structural details.

## Methodology

### Implementation of AlphaFold in PyTorch

In our study, we faced the challenge of AlphaFold’s unavailability for training code and its original implementation in JAX [3], which presents complexities in modification and testing. Inspired by the OpenFold model [2], we developed a custom version of AlphaFold using PyTorch library [21]. This approach not only allowed us to utilize the pre-trained weights of AlphaFold but also introduced significant flexibility in modifying the code as per our research needs.

One of the primary enhancements in our PyTorch-based AlphaFold implementation is the integration of checkpoints for optimal memory management. Given the extensive size of the AlphaFold network, these checkpoints are crucial for efficient memory usage, ensuring stable and effective processing even on large-scale data.

Furthermore, we leveraged the PyTorch-Lightning library [7], which significantly streamlined our workflow. PyTorch-Lightning abstracts and automates many routine tasks, enabling us to focus on the core aspects of our model. It facilitated effective training and easy distribution of computational workload across multiple GPUs and nodes. PyTorch-Lightning also brought additional advantages, such as simplified implementation of advanced optimization techniques and streamlined model validation processes. It enhanced our model’s reproducibility and scalability, allowing us to efficiently experiment with various configurations and settings.

### Dataset

We downloaded pMHC complex structures from the RCSB Protein Data Bank, selecting X-ray structures with a resolution finer than 3.5 Å. We excluded structures containing non-standard amino acids or a significant number of unresolved residues. Additionally, we limited our selection to samples with peptide lengths ranging from 8 to 11 amino acids. From each MHC protein only *α*1 and *α*2 domains were used. Our final dataset consisted of 944 structures from various species, including humans (Homo sapiens), mice (Mus musculus), and other species. The majority belongs to Homo sapiens (76%) and Mus musculus (mouse) (19%). We divided the dataset into three parts based on the release date, approximately a 60/20/20 split for training, validation, and testing. The training set (releases from 1992-10-15 to 2016-12-07) and the validation set (releases from 2016-12-07 to 2020-06-17) were used for fine-tuning and hyperparameter optimization, while the test set (releases from 2020-06-17 to 2023-08-23) was reserved solely for final comparison.

To prepare the necessary input for AlphaFold, we generated Multiple Sequence Alignments (MSAs) using MMseqs2, a software suite designed for fast and sensitive protein sequence searching. Our searches were conducted against the ColabFoldDB, which is a comprehensive and regularly updated database tailored for such analyses.

### Metrics

In protein structure prediction, the Root Mean Square Deviation (RMSD) measures the average atomic distance between a predicted protein structure and a reference, typically comparing backbone atoms. A prediction is considered accurate if the RMSD is below 2.0 Ångströms.

The Local Distance Difference Test (lDDT) [12] is another metric that evaluates the quality of a protein model at the local residue level. Unlike RMSD, lDDT is a superposition-free metric, meaning it does not require alignment of the structures and is insensitive to domain movements. It measures the local conformational similarity of each residue’s environment by comparing inter-atomic distances. The lDDT score ranges from 0 to 1, with higher values indicating better model quality.

AlphaFold’s predicted lDDT (plDDT) scores offer a valuable confidence measure for the predicted positions of residues within a protein structure. These scores are instrumental for researchers to gauge the reliability of specific regions within the predicted structural model, particularly when experimental structures are absent for validation. In our study, we have enhanced the original AlphaFold’s confidence predictions, enabling a more precise estimation of structure reliability.

Evaluating plDDT against lDDT can be done through comparison with known structures or through experimental validation. High plDDT values in regions that align with high lDDT scores from experimental data can indicate a successful prediction. To measure the quality of predicted lDDT scores, researchers often use statistical methods such as mean absolute error and the Pearson correlation coefficient.

In protein structure prediction, it is often necessary to focus on specific regions that are of functional significance. Accordingly, in our study, we computed the RMSD, lDDT, and plDDT scores with an emphasis on the peptide portions of the MHC-peptide complexes.

### Fine-tuning techniques

To refine the prediction capabilities of AlphaFold for MHC-peptide complexes, we started with the foundational AlphaFold multimer model v2.2. We explored several refinement techniques, focusing on both architectural modifications and training strategies.

#### Architectural Enhancements

Our initial approach involved augmenting the existing AlphaFold model by adding extra Evoformer blocks, ranging from 1 to 10, to the pre-existing 48. The design of AlphaFold, with its inherent residual connections, allows for such integrations without compromising existing functionalities, potentially enriching the model’s learning capacity.

#### Template Usage Optimization

Another significant adjustment pertained to the use of templates. While the standard AlphaFold architecture processes protein chains individually, we innovated by incorporating information regarding the interactions between protein chains and peptides. This modification is crucial for our focus on MHC-peptide interactions, aiming to capture the subtleties of these complex molecular interplays.

#### Focused Loss Function

Given our specific interest in the peptide structures, we introduced a novel weighting system within the loss function. This system assigns greater importance to the peptide residues over the protein residues, thus directing the model’s learning focus more towards the structural prediction of peptides.

#### Hyperparameter Tuning

On the training front, we experimented with various hyperparameters to optimize the model’s performance. This included testing different learning rate schedulers, such as CyclicLR, StepLR, and CosineAnnealingLR, with learning rates spanning from 0.001 to 0.000001. We also employed accumulate grad batches, serving as an analogue to batch size, to manage the model’s learning process more effectively. This broad range of hyperparameter experimentation allowed us to finely calibrate the model for optimal performance in predicting MHC-peptide structures.

The fine-tuning was conducted iteratively, evaluating each alteration to ensure that the modifications contributed positively to the model’s performance. The impact of these changes was quantitatively assessed using RMSD and lDDT scores as benchmarks for structural prediction accuracy.

## Results

### Optimal Parameter Settings: MHC-Fine Model

After extensive experimentation, we have identified an optimal set of parameters for our AlphaFold-based model, now termed ‘MHC-Fine’, tailored for predicting MHC-peptide complex structures. These parameters were determined as the most effective in balancing computational efficiency with predictive accuracy:

#### Additional Evoformer Blocks

We integrated two additional Evoformer blocks into the model. This choice was based on our observation that the structural similarity across our sample set did not necessitate a large increase in complexity. Two additional blocks provided the right balance for capturing the nuances of MHC-peptide interactions.

#### Template Utilization

The introduction of high-quality templates emerged as a crucial factor. Although the model was capable of learning independently of templates, their inclusion significantly accelerated the training convergence, underscoring their value in our fine-tuning process.

#### Hyperparameters

We employed the CosineAnnealingLR scheduler with a learning rate of 0.0003. This configuration facilitated a more dynamic adjustment of the learning rate, aiding in finer convergence. We set accumulate grad batches to 1, with the model being trained across 8 GPUs. This effectively meant a real batch size of 8, allowing for a more precise gradient update, which proved beneficial at this stage of the model’s training.

The MHC-Fine model, with these refined parameters, has shown the capability to outperform established benchmarks. Its single-model framework simplifies the prediction process while maintaining high accuracy, marking a significant step forward in computational immunology.

### Benchmarking Baseline Performances

To assess the baseline performance of AlphaFold on our test set prior to fine-tuning, we utilized all five multimer models from AlphaFold version 2.2. For each sample in the test set, we systematically evaluated the predicted structures generated by each model. The best result for each sample was selected based on the highest plDDT scores.

To ensure a comprehensive and fair comparison, we similarly evaluated the performance of the Pandora. We meticulously removed any instances where our test set overlapped with Pandora’s template database to avoid any potential bias from Pandora having prior knowledge of the structures in our test set. For each sample, the most probable structure predicted by Pandora was selected, using the molecular probability density function (molpdf) as the scoring function.

Employing this approach allowed us to establish a robust benchmark for the original performance of both AlphaFold and Pandora. This benchmark serves as a critical reference point against which we could measure the enhancements achieved through our fine-tuning process.

### Comparative Analysis

Our fine-tuning of AlphaFold led to a model with enhanced predictive performance for MHC-peptide complexes, showing a statistically significant reduction in median RMSD values compared to both the original AlphaFold and the homology-modeling-based Pandora approach. Figure 1 illustrates the distribution of Root Mean Square Deviation (RMSD) values for predicted MHC-peptide complex structures across three different computational methods: the original AlphaFold (median RMSD 1.44 Å), Pandora (1.27 Å) and our MHC-Fine model (0.65 Å). MHC-Fine model shows not only lower median RMSD but also a significantly narrower Interquartile Range (IQR), indicating higher accuracy and consistency in structure prediction. This suggests a closer approximation to the high-resolution crystal structures in our test dataset and indicates a marked improvement in the spatial accuracy of our model’s predictions.

**Fig. 1.**
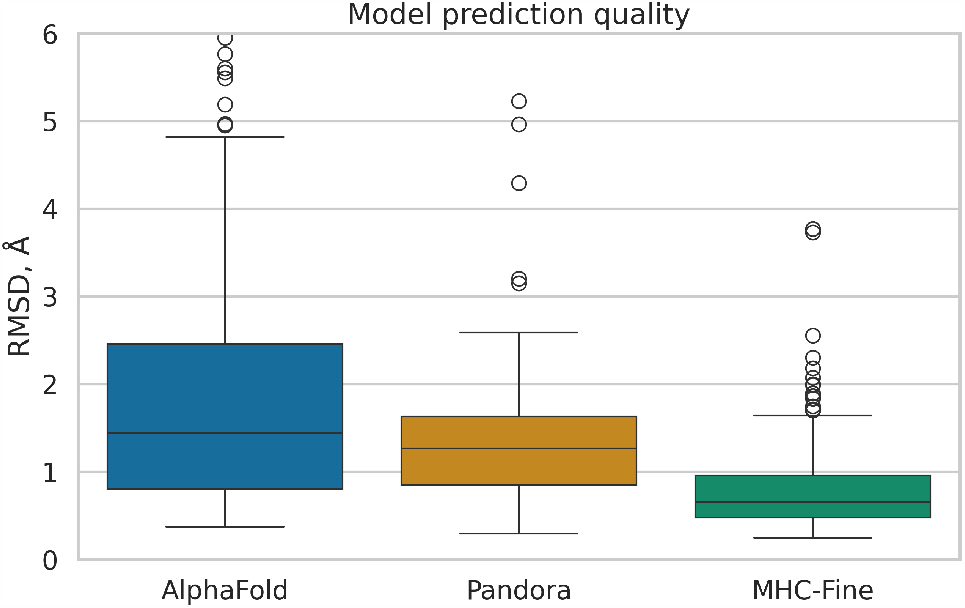
Comparative Analysis of Prediction Accuracy for MHC-Peptide Complexes

Our analysis is demonstrated through three case studies, each with varying prediction accuracies. Figure 2 illustrates these differences: (A) shows high accuracy with the fine-tuned AlphaFold, (B) depicts moderate accuracy for both models, and (C) highlights significant discrepancies in predictions.

**Fig. 2.**
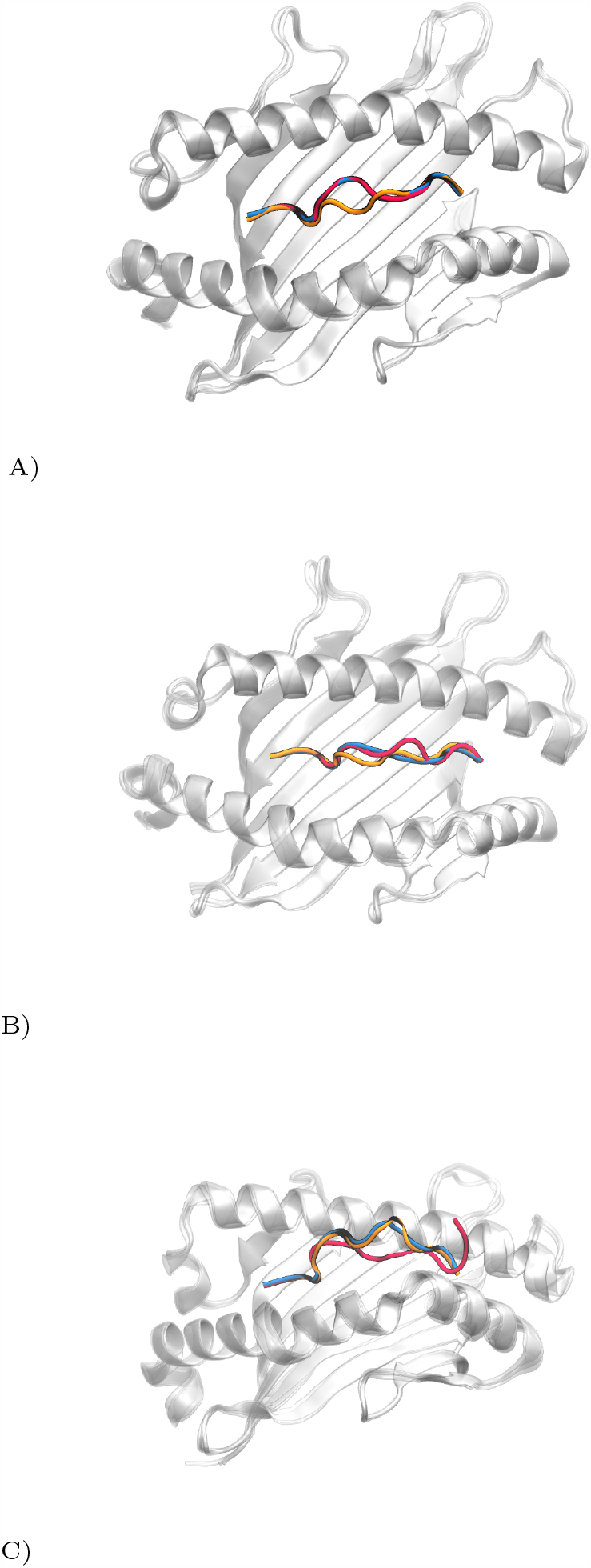
Comparative Visualization of pMHC Complex Prediction Accuracy. True peptide structure in red, Pandora model in orange, MHC-Fine (fine-tuned AlphaFold) in blue. (A) High precision of MHC-Fine (RMSD: 0.25 Å) versus Pandora (RMSD: 1.44 Å) for PDB ID: 6vb3; B)Moderate accuracy for both models (MHC-Fine RMSD: 1.23 Å; Pandora RMSD: 1.19 Å) for PDB ID: 7n2o; (C) Significant deviations in predictions (MHC-Fine RMSD: 3.73 Å; Pandora RMSD: 4.30 Å), indicating areas for enhancement, for PDB ID: 7mj7.

In addition to RMSD improvements, the fine-tuned model demonstrated enhanced lDDT scores, indicating higher accuracy compared to the original model. The mean absolute error was recorded at 3.0, and the Pearson correlation coefficient between actual lDDT scores and predicted lDDT values was 0.62, indicating a moderate positive relationship. Figure 3 illustrates that the MHC-Fine model produces a notably narrower error distribution for predicted lDDT values when compared to the original model. This more constrained spread signifies a higher consistency in the predictions, indicating that our fine-tuning process yields a more reliable and accurate measure of structural confidence across the majority of samples, with substantial deviations being considerably less frequent. Furthermore, Figure 4 shows that within our test dataset, a plDDT score greater than 90 is associated with an RMSD of less than 1.5 Å, and a plDDT score above 95 corresponds to an RMSD of less than 1.0 Å, underscoring the precision of our fine-tuned model in structural prediction.

**Fig. 3.**
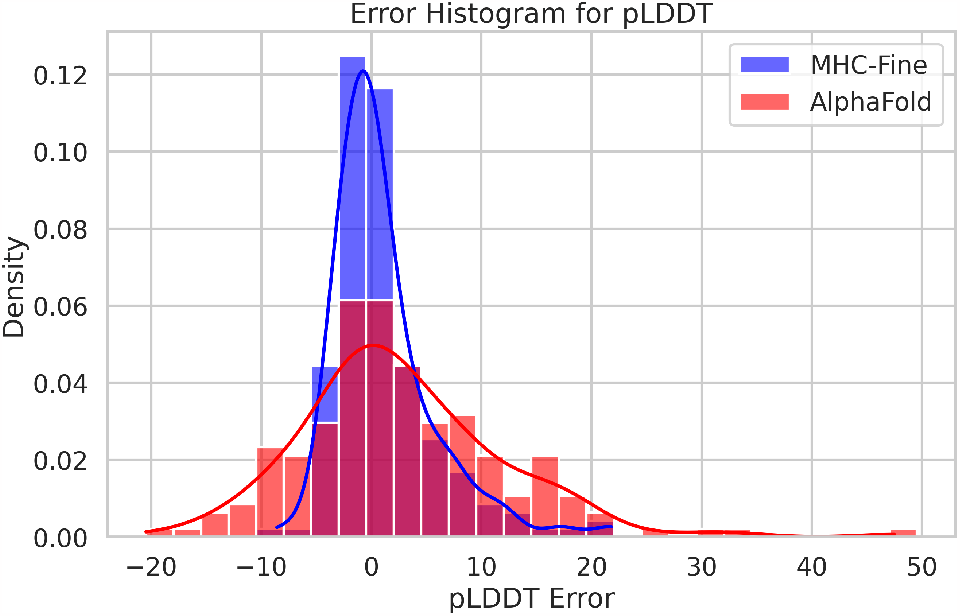
Error Distribution for Predicted lDDT Values. The red distribution represents the original AlphaFold model with a standard deviation of 9.2, while the blue distribution showcases our fine-tuned AlphaFold model with a reduced standard deviation of 4.6.

**Fig. 4.**
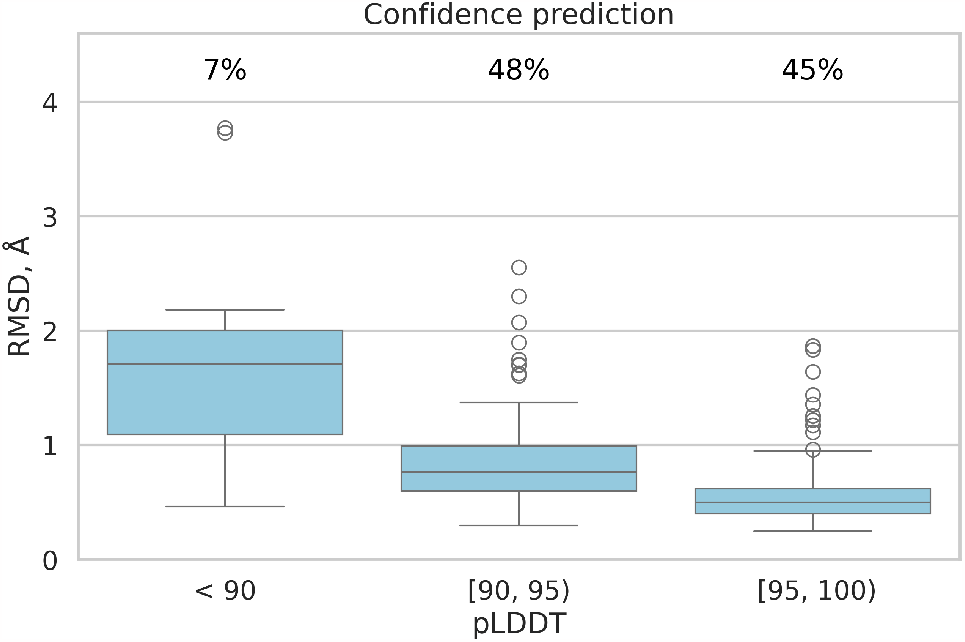
Confidence Prediction: All samples are grouped by their predicted Local Distance Difference Test (pLDDT) interval. The chart displays the percentage of samples falling within each specific pLDDT range.

## Conclusion

In conclusion, our study presents a refined AlphaFold model tailored for the intricate task of MHC-peptide complex structure prediction. By fine-tuning with high-resolution domain-specific data, we’ve achieved superior performance compared to both the original AlphaFold and traditional homology-modeling approaches. Our focused metrics on the peptide regions have yielded RMSD values indicative of high-precision predictions, while improvements in the plDDT scores reflect an enhanced confidence in the structural assessments provided by our model. These advancements hold promising implications for computational immunology, potentially expediting the discovery and design of novel therapeutics and vaccines.

## Code and Data Availability

The inference code and datasets utilized in this study are publicly available to facilitate further research via the following link: https://bitbucket.org/abc-group/mhc-fine/src/main/

## Acknowledgments

This work was supported in part by the National Institutes of Health grants RM1135136, R01GM140098; by the National Science Foundation grants DMS-1664644, DMS-2054251.

